# RNAseK is vital for epithelial proton pump activity in Drosophila melanogaster

**DOI:** 10.64898/2026.02.03.703209

**Authors:** C. Maurya, A.D. Gillen, S. Keenan, J.A.T. Dow

## Abstract

The transporting epithelial tissues comprising the *Drosophila melanogaster* alimentary and renal systems are known to share a common set of enriched genes sometimes referred to as the “epitheliome”, reflecting their shared transport functions. Core amongst these genes are the *vha* genes, which encode subunits of the large Vacuolar-type ATPase (V-ATPase) proton pump complex. However, many of the non-vha components of the epitheliome remain broadly uncharacterised. Here, we explore the role of RNAseK, a gene identified during unbiased epithelial screens in *Drosophila* whose function within insects is not yet known, though evidence from mammalian systems suggest a role in supporting proton pump activity. We demonstrate computationally that RNAseK is strongly conserved across evolutionary history, and that expression is regulated by the highly epithelial-specific dCLEAR motif. Seeking to understand why epithelial expression is so emphasised, we have assayed the effects of RNAseK knockdown in different epithelia throughout the fly. Across hindgut, midgut, and Malpighian tubules, we note profound defects in gross tissue morphology, transport activity, and fly survival. Mechanistically, our findings that RNAseK may co-localise with the V-ATPase complex, and that V-ATPase inhibition phenocopies RNAseK knockdown, suggest that RNAseK is a critical component of the proton transport axis across *Drosophila* tissues.

**Summary statement:** RNAseK is enriched throughout, and required in, *Drosophila melanogaster* epithelial tissues. Molecular evidence and evolutionary inferences suggest this is due to a role in the proton transport axis.

## Introduction

The fruit fly *Drosophila melanogaster* provides a prime opportunity for interrogating tissue function in the context of a rich experimental genetic framework. Aside from providing direct insights into fly biology, *Drosophila* tissues share a high degree of functional homology with human tissues, so studying tissue function in the fly can be an efficient strategy to study human disease (Verheyen, 2022; Pandey and Nichols, 2011). *Drosophila* research can also provide a window into insect biology as possibly the most well-studied species within *Hexapoda*. As such, understanding tissue function within the fruit fly, especially in response to experimental pesticides, provides an intuitive avenue to predicting functionality within pest species (Ali et al., 2022; Denecke et al., 2022; Nainu et al., 2022).

Tissue function is determined by tissue-specific gene expression programs, which specify cell fate, functionality and response to stimuli (Chintapalli et al., 2013; McDonough-Goldstein et al., 2021; Regadas et al., 2021). This is also true for cell-type function, controlled by cell-specific expression patterning. Notably, it can also prove beneficial to combine tissues into functional sets for analysis. For example, investigating genes expressed throughout the female (McDonough-Goldstein et al., 2021) and male (Takemori and Yamamoto, 2009) reproductive tissues has revealed genes common to all tissues which are crucial to sex-specific biology.

Similarly, *Drosophila’s* alimentary canal, composed of fore-, mid-, and hindgut, as well as salivary glands and Malpighian tubules, lends itself to this form of functional grouping. As a whole, these tissues play an important role in nutrient transport, either in fluid secretion (salivary gland (Maruyama and Andrew., 2012) and Malpighian tubules (Beyenbach, Skaer and Dow, 2010) or nutrient absorption (midgut and hindgut; (Colombani and Andersen, 2020)). For this reason, these tissues may be referred to collectively as the “transporting epithelia”. Previous efforts into identifying key genetic determinants of the transporting epithelia, termed the “epitheliome”, have identified several genes whose expression appears to be closely linked to epithelial specification (Chintapalli et al., 2013; Gillen et al., 2024).

Core amongst epitheliome components are several subunits of the Vacuolar Type ATPase (V-ATPase) holoenzyme. V-ATPases are membrane bound proton pumps composed of 15 subunits, which can be divided into a cytoplasmic V1 region and a membrane-bound V0 region (Dow et al., 1997). V-ATPases are critical for many aspects of cell biology, including maintenance of electrostatic gradients (Dow et al., 1997; D’Silva, Donini and O’Donnell); and maintaining appropriate pH balance in cells and vesicles (Dulac et al., 2021; Yan, Denef and Schupbach, 2009). As a result of these diverse activities, V-ATPase activity is ubiquitous, and not limited to epithelial tissues (Dulac et al., 2021, Jain et al., 2024).

That said, the 15 canonical V-ATPase subunits are encoded by 33 Vha genes within Drosophila, allowing numerous theoretically possible configurations of the overall holoenzyme structure (19). Given this fact, it is striking that epithelial tissues consistently and preferentially express 15 Vha genes comprising a distinct V-ATPase holoenzyme from non-epithelial tissues (Chintapalli, 2013; Gillen et al, 2024; Chintapalli, Wang and Dow, 2007), an expression profile which seems to be co-ordinated by the transcription factor MITF (Bouché et al., 2016).

Intriguingly, though, there is evidence that this view of epithelial V-ATPase function remains incomplete. In particular, evidence from other species has recently uncovered roles for homologues of RNAseK in supporting the function of V-ATPase (Makar et al, 2024; Perreira et al, 2015). Hypothesis-free analysis of *Drosophila* epitheliome components has also identified RNaseK as extremely closely linked to epithelial function (Gillen et al., 2024). Based on these data, we hypothesised that RNAseK may act as a *de facto* part of the epithelial-specific V-ATPase holoenzyme within *Drosophila*, thereby playing a core part of pan-epithelial function. Here, we provide an overview of transcriptomic and evolutionary data supporting this assertion; demonstrate a role for RNAseK throughout Drosophila epithelia; and show evidence linking this role to V-ATPase function in an epithelial, rather than an intracellular, context.

## Materials and Methods

### Data Utilisation

*Drosophila melanogaster* annotation information were obtained from FlyBase (release FB2025_03). Specifically, a list *Drosophila* transcription factors were compiled based on FlyBase gene group FBgg0000745, and gene identifier mapping information was derived from fbgn_fbtr_fbpp_expanded_fb_2025_03.tsv.gz.

*Drosophila melanogaster* transcription factor binding motifs were obtained by combining the interactome reported by Shokri et al (2019) with those available through CisBP (Weirauch et al., 2014).

### Regulatory Motif Analysis

Analysis of known regulatory motifs shared within a gene set was carried out using i-cisTarget (Imrichová et al., 2015) (https://gbiomed.kuleuven.be/apps/lcb/i-cisTarget/index.php). The settings were as follows: Full motif collection; i-cisTarget motif database v 6.0; for Drosophila melanogaster (dm6) with gene annotation FlyBase 6.02. The search region was defined as 5 kb upstream of each gene’s transcriptional start site to the end of the transcript, with an 0.4 minimum fraction of overlap with identified motifs. Normalised Enrichment Score thresholding was set to 3 (the default), based on enrichment calculated within each motif data separately and an ROC threshold of 0.01 for area-under-curve analysis.

De novo motif identification was carried out using FindMotifs, a part of the Hypergeometric Optimization of Motif EnRichment (HOMER) suite (Heinz et al., 2010) (http://homer.ucsd.edu/homer/, version v5.1, 7-16-2024). Enrichment of sequence motifs over the HOMER Drosophila melanogaster promoter set. The search region was defined as 2 kb upstream of each gene’s transcriptional start site to 2 kb downstream – the maximal search region permitted within HOMER. Other settings for this analysis were as follows: Repeats were masked; Motifs up to 14 bp searched; all other settings at default.

Additionally, new software was developed to check for motif occurrence and enrichment at the same time. Using BLAMM for motif searching (Fostier, 2020), all genes (+/- a defined window size) are checked for occurrences of a given motif. Specific genomic co-ordinates for each occurrence are returned and, if multiple genes of interest are specified, motif frequency is compared with general motif occurrence rates using χ^2^ testing. This software is freely available at https://github.com/AndrewDGillen/MotifScanning and on Figshare at 10.6084/m9.figshare.31037749

### Multiple Sequence Alignment

Multiple Sequence Alignment was run using Muscle5 (Edgar, 2022) via the EBI online portal (https://www.ebi.ac.uk/jdispatcher/msa/muscle5) using default settings. This was run three times, to analyse conservation separately for RNAseK Protein, cDNA, and gene sequences.

RNAseK orthologues were identified using OrthoDB (Tegenfeldt et al., 2025). Orthologues were as follows: *Anopheles gambiae* ENSAGBG00005004689; *Aedes aegypti* AAEL002492; *Schistocerca gregaria* XM_049992438.1*; Bombyx mori* LOC732881 and LOC101747017; *Tribolium castaneum* TC008797 and TC006752; *Adalia bipunctata* ENSABPT00005007386.1; *Apis melliflera* LOC725572; *Danio rerio* ENSDARG00000104458 and ENSDARG00000069461; *Mus musculus* ENSMUSG00000093989; *Homo sapiens* ribonucleaseK

### Fly stocks and husbandry

Drosophila lines of the desired genotypes were maintained on standard Glasgow Drosophila diet (Dornan et al., 2023) under controlled conditions of 45–55% relative humidity, a 12-hour light/dark cycle, and a temperature of 24 ± 1 °C in a SANYO incubator unless otherwise specified. For GAL4/UAS system experiments with the tub Gal80^TS^ transgene, parental (control) and F1 (experimental) flies were initially reared at the permissive temperature of 18 °C. After eclosion, experimental F1 progeny were shifted to the restrictive temperature of 29 °C to inactivate Gal80^TS^ and permit gene expression; simultaneously, control flies were also transferred to 29 °C to ensure age-matched comparisons.

The following fly lines were acquired from the Bloomington Drosophila Stock Centre (Bloomington, IN, USA):

y v; P{y[+t7.7]=CaryP}attP2 (BL #36303); y sc[] v sev; P{y[+t7.7] v[+t1.8]=TRiP.HMS01529}attP2/TM3, Sb (BL #35779); y w[]; PBac{y[+mDint2] w[+mC]=UAS-RNASEK.HA}VK00033 (BL #94316); w; P{w[+mC]=PTT-GA}VhaSFD[G00259]/CyO (BL #6840); P{w[+m*]=sens-GAL4.B}1, y w[] (BL #42740).

Myo31 df Gal4 ; tub Gal80^TS^ (Adam Dobson Lab, University of Glasgow, Scotland, UK); Da Gal4; tub Gal80^TS^ (Ilan Davies Lab, University of Glasgow, Scotland, UK); Irk-2 Gal4/Cyo; UAS-GFP/TM6c (B. Denholm Lab, University of Edinburgh, Scotland, UK) were obtained from collaborative laboratories.

Other experimental lines: w*; CapaR-GAL4 (Terhzaz et al., 2012), w*; UAS mCD8::GFP; CapaR-GAL4’, ‘C-724 GAL4 (Sözen et al., 1997), and introgressed line w; C-724 GAL4; UAS-mGFP, (Dornan et al., 2023) were generated and maintained in-house.

### Immunocytochemistry and Phalloidin staining

Dissected tissues from 9-day old adult Drosophila were collected in Schneider’s Drosophila medium (Gibco, Thermo Fisher Scientific, Cat #20012-019) at room temperature, then fixed in freshly prepared 4% PFA in 1× phosphate-buffered saline (PBS) for 20 minutes to ensure cellular structure preservation. After fixation, the tissues were permeabilised by three successive 15-minute washes with PBS containing 0.1% Triton X-100, PBST, (Sigma-Aldrich, Catalog No. 93443-100ML, 10% stock in H□O). For blocking, tissues were incubated for two hours at room temperature in a solution of 5% Normal Goat Serum and 0.5% Triton X-100 in 1× PBS (Gibco, Ref 20012-019).

For immunostaining, tissues were incubated overnight at 4 °C in blocking buffer containing mouse monoclonal anti-Dlg1 (1:50, 4F3, DSHB) and rabbit polyclonal anti-RNAseK (1:1000, Geneosphere Biotechnologies). The RNAseK antibody was custom-generated in host rabbit against the peptide Heptan ‘N’-EDLPLEEEYHSLEDC, coupled to keyhole limpet hemocyanin (KLH), and affinity purified with ELISA verification (absorptions for positives were >2-fold over pre-immune serum).

Following the primary antibody step, tissues were washed three times for 15 minutes each in 0.1% PBST. For secondary detection, Alexa Fluor 546-conjugated anti-rabbit IgG (cat. no. A-11035, Thermo Fisher Scientific) and Alexa Fluor 633-conjugated anti-mouse IgG (cat. no. A-21052, Thermo Fisher Scientific) were each diluted 1:500 and incubated with samples for two hours at 37 °C. After secondary incubation, tissues were washed and then treated with Dapi (1 μg/ml, Thermo Fisher Scientific, Cat No. D21490) prior to mounting on Polysine slides (VWR International, Cat # 631-0107) with antifade medium, followed by imaging using a Zeiss LSM 880 confocal microscope (Carl Zeiss).

For phalloidin staining, tissues were dissected, fixed, and permeabilised in the same way but incubated with Phalloidin–Atto 550 (Sigma-Aldrich, Cat No. 19083) at 1:1000 dilution for 15 minutes at room temperature to stain F-actin. After staining, tissues were washed three times in 0.1% PBST, followed by incubation in Dapi (1 μg/ml) for 10 minutes and three additional washes with 0.1% PBST. These samples were also mounted on Polysine slides with antifade medium as above and imaged on the Zeiss LSM 880 confocal microscope. Confocal image stacks were analysed with ImageJ (ImageJ 2.16.0), and representative z-projections were prepared for figure assembly in Microsoft PowerPoint (Version 2405)

### Stellate cell counting

For quantification of stellate cells in control and RNAseK downregulated flies, C-724 transgenic flies recombined with mGFP were crossed with UAS control and RNAseK knockdown flies. The Malpighian tubules were dissected from 9-day old adult female flies in 1× PBS and fixed in 4% PFA for 10 minutes at room temperature. Fixed tubules were mounted on glass slides using antifade mounting medium, and stellate cells were manually counted under a fluorescence microscope. Counts from anterior and posterior tubules were recorded separately. The raw data were analysed using GraphPad Prism 10 (Version 10.2.3), and statistical significance was determined using multiple t-test (nonparametric).

### RNA isolation, cDNA synthesis and qRT-PCR

RNA was isolated from 50 pairs of freshly dissected adult Drosophila tubules using QIAzol Lysis Reagent (QIAGEN Sciences USA, 50 ml) followed by purification with the miRNeasy Mini Kit (Cat # 217004, QIAGEN Sciences USA). The final RNA prep was resuspended in nuclease-free dH□O provided with the kit. For cDNA synthesis, 300 ng of total RNA was used in a SuperScript II RT reaction (Cat # 18064-022, Thermo Fisher Scientific), according to the manufacturer’s instructions. qRT-PCR was carried out using the QuantiNova SYBR Green PCR Kit (Cat # 208054, 500 reactions, QIAGEN) on a StepOnePlus Real-Time PCR system (Applied Biosystems), with gene-specific primers for Rpl32 (Rpl32—F: TTTCGCTTCTGGTTTCCGG; Rpl32—R: CTTGCGCCATTTGTGCGA), dPrestin (dPrestin-F: AGTTTGCCAAGGTGCGTTC; dPrestin-R: CAGGAGTACGGCTTCAGTCC), SPoCk (SPoCk-F: GCATTCGTCATTTTTGCTGCG; SPoCk-R: AGGGTAAGGCCCGTCTGTT), Dh44-R2 (Dh44-R2—F: GGTCGATAGCCTGGATGACG; Dh44-R2—R: GAATCGAACGAGCTGGGACA), Crystallin (Crys-F: CAGCACCAGATTTGAGCGTC; Crys—R: GTGAGAGAGGGGCGAAACTC), RNase-K (RNase-K—F: TGTGGCCCGAAACTATCACTT; RNase-K—R: GCAGTTGTAGGCATTCTGATTGT), Vha68-2 (Vha68-2-F: GAACGCGAAGACACAGCAAA; Vha68-2-R: CAGTTACGCCAGAGGTCTCC), (Integrated DNA Technologies, Belgium).

Primer specificity was confirmed by running PCR products on agarose gels using cDNA as template -with gDNA used as a negative control on the same gel - and by checking for single melt curve peaks in qPCR. For each 96-well plate (Cat # 4346907), 5-fold dilution series standard curves were generated in duplicate alongside sample amplifications to ensure amplification met the MIQE standard 90-110% efficiency threshold (Bustin et al., 2009). Results from qRT-PCR analysis were normalised to Rpl32 expression, providing the gene/Rpl32 expression ratio per sample. Final data were plotted using GraphPad Prism 10 (Version 10.2.3).

### Starvation and Desiccation assays using DAM5H Monitors

All experiments were performed using the DAM5H Drosophila Activity Monitor (TriKinetics Inc., USA) under standard laboratory conditions (24 ± 1 °C; ambient humidity maintained within the DAM incubator). Flies aged 9-days (both sexes) were briefly anesthetised and loaded individually into 65 mm glass tubes. In the starvation assay, tubes were prepared with approximately 1 cm of 1% agar at one end for humidity, sealed with a rubber plug, and flies were introduced from the opposite end, which was closed with cotton to maintain aeration; activity was then monitored in the DAM5H for up to 6-days based on optimised female survival. For desiccation, flies were placed in empty tubes sealed with cotton at both ends and monitored for 4-days. After completion, data were extracted from the DAM system, compiled in Microsoft Excel (Version 2405), and analysed in GraphPad Prism 10 (Version 10.2.3) to identify outliers and compare groups using an unpaired non-parametric Mann-Whitney test for rank comparisons. All experiments were carried out in biological triplicate, and final results were drawn from compiled replicates.

### Midgut pH assays and Gut length measurement

m-Cresol purple (Cat # 857890-5G, Sigma-Aldrich) indicator dye was mixed into melted Standard Glasgow Drosophila diet containing 1% Kaolin (Cat # K7375-500G, Sigma-Aldrich), cooled to 60 °C (final concentration 0.3% w/v), then swirled and allowed to reach room temperature.

Adult female flies, 9-day old, of the appropriate genotype were placed on the dye-mixed food for overnight feeding. The next day, midguts were dissected in Schneider’s Drosophila medium (Cat # 21720-024, Thermo Fisher Scientific), and immediate micrographs were taken with a Zeiss Stemi 508 stereomicroscope equipped with an Axiocam 105 color camera, as pH remains stable for only a few minutes after dissection. Images were acquired using Zeiss Zen 3.9 software.

For gut length measurements, individual gut images were opened in Zeiss Zen 3.9 software and the length was traced and measured using the spline tool. Recorded lengths were analysed in GraphPad Prism 10 (Version 10.2.3) using an unpaired Mann-Whitney test for rank comparisons.

### Ramsay fluid secretion assay

Fluid secretion assays using Drosophila Malpighian tubules (MTs) were performed following established protocols (Halberg et al., 2015; Dow et al., 1994). Anterior tubules were dissected in ice-cold Schneider’s medium and placed in a 9 µl drop containing a 1:1 mix of Schneider’s medium and Drosophila saline with trace amaranth dye (cat no. A1016; Sigma-Aldrich). After acclimatisation, baseline secretion rates were measured for each tubule at 10 min intervals over a 30 min period. Subsequently, 1 µl of Capa neuropeptide (10^−6^ M; Solasta Bio, Scotland, UK) was added to the drop, and fluid secretion rates were measured every 10 min for an additional 30 min. A diuretic effect was confirmed if secretion increased after Capa application compared to basal levels.

This experiment was performed using Malpighian tubules from 9-days old female flies in three independent biological replicates. For final analysis, data from all replicates were combined, and statistical significance was determined using two-way ANOVA with the Geisser-Greenhouse correction, a family-wise alpha threshold, and comparison with a 95% confidence interval. Graphs were generated in GraphPad Prism 10 (Version 10.2.3).

### Bafilomycin feeding and Gut analysis

9-day old Canton-S control flies (10 per group) were used for experiment, during which they were maintained on 5% sucrose solution for the entire 9-day period. For the treatment group, flies were placed in separate vials containing 200 µL of 5% sucrose solution supplemented with 50 µM bafilomycin A1 (Catalog No: 54645, Cell Signaling Technologies). Flies were sorted into control and treatment groups since day 1 of age and reared until day 9. The feeding solution was replaced daily, and all vials were kept in a dark chamber throughout the assay to prevent light-induced degradation of bafilomycin A1 and to ensure its stability.

To analyse gut pH partitioning following treatment, flies were given overnight access to food containing m-Cresol purple dye (Catalog No: 857890-5G, Sigma-Aldrich) and 1% kaolin (Catalog No: K7375-500G, Sigma-Aldrich), following the midgut pH measurement protocol. Immediately after feeding, midguts were dissected and imaged using a Zeiss Stemi 508 stereomicroscope with Axiocam 105 color camera, and images as well as gut length measurements were processed using Zeiss Zen 3.9 software for both bafilomycin-treated and control flies.

### Fly weight and Water content measurements

To measure wet body weight, 20 9-day old Drosophila from each replicate (n = 9; consisting of three biological replicates with three technical replicates per biological replicate, and both genders included) were anesthetised and transferred into pre-weighed 1.5 mL tubes (Cat # 616 201, Greiner Bio-One). The flies were weighed immediately after transfer using a GR-202 analytical balance (A&D Instruments, Abingdon, UK). For dry body weight measurement, flies were first killed by freezing at –80 °C for 20 minutes, then dried at 65 °C for 72 hours in a Hybridiser HB-1D (Techne). After reaching room temperature post-drying, flies were weighed again using the same analytical balance. Water content was calculated based on the difference between wet and dry weights. Percentage water content, along with all weight data, was statistically analysed using an unpaired Mann-Whitney test for rank comparisons in GraphPad Prism 10 (Version 10.2.3).

### SMURF intestinal permeability assay

For assessing intestinal barrier integrity, the Smurf assay was performed following the protocol adapted from Bio-protocol (Martins et al., 2018). Standard melted fly food was cooled to approximately 60 °C, and FD&C Blue Dye #1 (Cat # 861146, Sigma-Aldrich) was added at a final concentration of 2.5% w/v (2.5 g per 100 ml). The food was gently mixed, swirled to ensure even dye distribution, and allowed to cool fully to room temperature. Adult male and female flies of the desired genotypes were transferred to freshly prepared dyed food and maintained for 14 hours of feeding at standard laboratory conditions.

After overnight exposure, flies were transferred to fresh, non-dyed standard media for immediate visualization. Scoring was performed via visual inspection for blue coloration throughout the body - the so-called Smurf phenotype - which indicates intestinal barrier dysfunction and leakage of the non-absorbable blue dye beyond the digestive tract. The flies were classified into Non-Smurf, Uncertain, Light Smurf, and Smurf categories, and the proportions were converted to percentages. The graph was plotted using GraphPad Prism 10 (Version 10.2.3).

### Drosophila feeding assay (pipette-tip)

To conduct the Drosophila feeding assay, individual 200 µL pipette tips (Cat # S1120-8810, TipOne Pipette Tips, Starlab) were first weighed using a GR-202 analytical balance (A&D Instruments, Abingdon, UK) to record their initial empty weight (W1). Each tip was then filled with 30 µL of freshly prepared 5% sucrose solution, and the weight was recorded again (W2). Ten adult flies (9-day old) were introduced into each assay vial, and a loaded pipette tip was inserted vertically to ensure flies could access the sucrose solution freely. The vials were maintained at 25 °C in a SANYO incubator for 24 hours, allowing the flies to feed. After 24 hours, the pipette tip was removed and weighed to determine the final weight (W3). The volume of sucrose consumed was calculated by subtracting W3 from W2, converting this weight change to microliters. Feeding data were analysed statistically using an unpaired t-test Mann-Whitney test, in GraphPad Prism 10 (Version 10.2.3).

### Embryo fixation and visualisation

Embryos (20-22 hrs) were dechorionated by immersing them in freshly prepared 50% sodium hypochlorite (bleach) solution for 2 minutes with gentle agitation, effectively removing the outer chorion layer. After dechorionation, embryos were thoroughly washed with double distilled water to eliminate any remaining bleach. For permeabilization and fixation, embryos were vigorously shaken in a 1:1 mixture of 4% PFA fixative and n-heptane (Catalog No: GPS9716, apc pure) for 30 seconds, ensuring the fixative could penetrate the vitelline membrane efficiently. The embryos were then washed three times with 0.1% PBST, each wash lasting 3 minutes to remove any residual fixatives and permeabilizing agents. Finally, embryos were mounted onto slides and visualised under polarised light using a Zeiss Stemi 508 stereomicroscope equipped with an Axiocam 105 color camera. Micrographs were acquired using Zeiss Zen 3.9 software for subsequent analysis.

### Alizarin Red Staining

Malpighian tubules from 9-day old flies of the desired genotypes were dissected in 1× PBS solution and immediately fixed in 4% PFA for 15 minutes to preserve tissue architecture. The fixed tubules were then rinsed three times for 5 minutes each in 0.1% PBST to ensure thorough permeabilization. Staining was performed by incubating the tubules in Alizarin Red S (Catalog No: A5533-25G, Sigma-Aldrich) at a concentration of 20 mg/ml (filtered twice using Whatman^TM^ Cat No 1001 125) dissolved in 0.1% PBST for 4 minutes.

After staining, the tubules were destained and rinsed three times in 0.1% PBST for 5 minutes each on a shaker to remove excess dye. The stained tubules were then mounted and imaged under transmitted polarised light using a Zeiss Stemi 508 stereomicroscope equipped with an Axiocam 105 color camera. Images were acquired and analysed using Zeiss Zen 3.9 software for presentation.

### Abdominal width determination

Flies of the desired genotype and age were positioned laterally for imaging using a Zeiss Stemi 508 stereomicroscope equipped with an Axiocam 105 color camera. The acquired image was opened in Zeiss Zen 3.9 software, where abdominal width was measured. The resulting values were statistically analysed using an unpaired Mann-Whitney test for rank comparisons, in GraphPad Prism 10 (Version 10.2.3), allowing for robust comparison of abdominal size across genotypes or experimental groups.

### RNAseK follows epithelial transcriptional programs

Hypothesis-free analysis of large, publicly available data have demonstrated that RNAseK broadly follows a pattern of enrichment across epithelial tissues within *Drosophila melanogaster* (Cohen et al., 2020). Strikingly, this closely mirrors the evidence supporting epithelial enrichment of V-ATPase subunits (**Fig. 1A**).

The epithelial pattern of *vha* genes contributing to the eptihelial V-ATPase is maintained by the transcriptional regulator MITF, via a motif referred to as dCLEAR (Bouché et al., 2016). To investigate whether this motif could also be responsible for co-ordinating epithelial RNAseK, we developed software to study the motifs around a given gene (https://github.com/AndrewDGillen/MotifScanning).

Using this software, we identified two putative dCLEAR-like motifs within two kilobases (kB) of the *RNAseK* gene, one overlapping the *RNAseK* transcriptional start site (2R:4188117-4188127) and one within 2 kB of the *RNAsek* end site (2R:4184962-4184972) (**Fig. 1B**).

To demonstrate whether the dCLEAR motifs identified in *RNAseK* and the epithelial *vha* genes, we used our software to analyse motif occurrence across these 16 genes, and compared the rate of dCLEAR detection with the rest of the known *Drosophila melanogaster* genes. As previously described (Bouché et al., 2016), Drosophila Mitf regulates the V-ATPase and the lysosomal-autophagic pathway. *Autophagy*, **12**, 484-498. all 15 epithelial *vha* genes are within 2 kB of a dCLEAR motif. On the other hand, this is true of only 43% of genes globally – a difference which χ2 testing confirmed is statistically significant (adj p<0.0001).

Independently of our targeted analysis, we ran *de novo* motif detection software on our *RNAseK* plus *vha* gene set. Both i-cisTarget and HOMER motif detection returned very similar motifs associated with this gene set (**Fig. 1C**), which bore striking similarity to the known dCLEAR motif. As such, there is strong evidence that RNAseK, as with the epithelial V-ATPase subunits, is regulated by MITF within *Drosophila* epithelial tissues.

### RNAseK is strongly conserved across evolutionary history

Given this evidence of a shared transcriptional program; and the wealth of evidence that RNAseK is a V-ATPase binding partner in non-fruit fly species (Makar et al., 2024; Perreira et al., 2015), we hypothesised that this activity could be conserved across evolutionary history. As such, we sought to explore conservation of RNAseK homologues as defined by OrthoDB (Tegenfeldt et al., 2025) from *Drosophila,* non-fly insects, and other model organisms (**Fig. 2**).

The structure of RNAseK as predicted by AlphaFold (Jumper, 2021) is shown in **Fig. 2A**, along with a diagrammatic representation of the domains contained within the protein, based on SMART predictions (Letunic, Khedkar and Bork, 2021) (**Fig. 2B**). Critically, this predicts three major structural domains – two transmembrane domains and a low-complexity region.

Within this context, the conservation of RNAseK functionality can be more fully understood. Species of interest were aligned to *Drosophila melanogaster* RNAseK protein sequence, and conservation recorded across the sequence (**Fig. 2C**). Notably, conservation is high around the end of the first transmembrane domain; the low-complexity region and the entirety of the second transmembrane domain; and the end of the protein. Thus, it is entirely conceivable that the function of these domains has been conserved despite enormous evolutionary distance, implying a critical function.

The conservation of RNAseK protein, cDNA, and gene sequence globally was also measured for all orthologues via Multiple Sequence Alignment (**Fig. 2D**). Conservation was generally very high amongst other insect species, though protein conservation was higher than cDNA conservation, which was in turn higher than gene conservation. This pattern is the reverse of what is typical across species (Shih et al 2015), so may imply that the structure of RNAseK protein is under intense selective pressure for conservation.

Interestingly, though, the sequence divergence of RNAseK is not one-to-one with the course evolutionary divergence – some homologues show particularly stong or weak conservation. For example, *Schistocerca gregaria* shows high similarity to *Drosophila* RNAseK despite being from a wildly divergent taxonomic Order; whilst the mosquito *Aedes aegypti* shows very limited conservation despite also being a Dipteran species. The evolutionary relationships between studied species, along with phylogenetic trees based on sequence identities, are shown in **Fig. S1.**

### RNAseK loss disrupts embryonic tubulogenesis and adult feeding ability

Previous functional analyses of vacuolar ATPase (V□ATPase) subunits such as vha55 established their indispensable role in epithelial polarity and tubular morphogenesis in Drosophila (Davies et al., 1996; Dow and Davies, 2001; Allan et al., 2005), prompting assessment of whether the V□ATPase□associated factor RNAseK (Perreira et al., 2015) is similarly essential.

Embryos maintained at 25 °C and imaged 20-22 hours after egg laying formed well□organised Malpighian tubules under polarised light in controls (**Fig. 3A**) (Myat, 2005; Denholm, 2023; Cohen et al., 2020), whereas RNAseK deficiency caused complete absence of tubule formation (**Fig. 3B**), reminiscent of Vha55 or epithelial□polarity mutants showing tubular collapse during embryogenesis (Davies et al., 1996; Wieczorek et al., 2009).

In adults, flies grown at 18 °C until eclosion and shifted to 29 °C for conditional RNAseK knockdown showed normal body (**Fig. 3C-D**) and tubule morphology (**Fig. 3E-F**) and unchanged abdominal width (**Fig. 3G; Fig. S2, C**), indicating preserved renal structure. However, feeding assays revealed significantly reduced nutrient intake in RNAseK downregulated adults (**Fig. 3H; Fig. S2, E**), paralleling loss of proton□pumping efficiency seen in V□ATPase mutants that compromise secretory granule maturation in salivary glands (Farkaš et al., 2015).

### RNAseK regulates multiple critical midgut functions

Proton transport by the V-ATPase complex establishes the distinctive midgut pH gradient in Drosophila, vital for digestion and copper-cell physiology (Shanbhag and Tripathi, 2005; Strand and Micchelli, 2011; Overend et al., 2016). Control females displayed a typical midgut acidification pattern (**Fig. 2A**; **Fig. S3, B)**, as reported by Dubreuil (2004); while targeted RNAseK knockdown resulted in loss of the characteristic acidic copper-cell zone (**Fig. 4B; Fig. S3, B**). This phenocopies V-ATPase subunit vha100-2, vha100-4, and vha55 loss-of-function (Overend et al., 2016), confirming RNAseK dependence for lumenal proton gradient formation.

Treatment with Bafilomycin A1, a V-ATPase inhibitor, recapitulated the RNAseK RNAi phenotype disrupted acidic colouring, abdominal bloating, and enlarged gut diameter (**Fig. 4D-H**; **Fig. S3, F**)—consistent with failed V_0_ domain H□translocation (Mauvezin and Neufeld, 2015; Wang et al., 2021). Interestingly, targeted RNAseK downregulation in the hindgut using Irk-2 Gal4 resulted in remarkable localised swelling presumably functional blockage, likely impacting water excretion and causing the gut to swell **(Fig. S3, G**).

Assays investigating pH gradients in this context revealed a complete loss of the acidic (red) region, which was replaced by a neutral (yellow) profile in those cells (**Fig. S3, G**). This indicates a pronounced disruption of normal pH partitioning, possibly due to neutralization from abnormal water reflux or unidentified mechanisms. A similar phenotype was seen when the V-ATPase subunit Vha-SFD was mutated, also resulting in loss of the characteristic red acidic region from the midgut and the emergence of a neutral pH, suggesting inactivation of the proton pump (**Fig. S3, H**). These findings reinforce that both RNAseK and classic V-ATPase subunits are essential for compartmentalization of gut pH gradients, and their loss leads to consistent disruption of acid regions through pump inactivation, as supported by Overend et al., 2016 and Dubreuil., 2004.

Given the crucial physiological implications, we were particularly interested in the role of RNAseK in stress response. Consistently, stress assays showed markedly reduced resistance (**Fig. 4I**) and shortened gut length (**Fig. 4J**) when RNAseK was genetically downregulated, indicating impaired epithelial maintenance associated with V-ATPase disorganization (Perreira et al., 2015; Overend et al., 2016). Phalloidin staining upon RNAseK knockdown showed disrupted cortical F-actin and cytoskeletal collapse (**Fig. 4L-K**), consistent with V-ATPase–actin coupling via the WASH/Arp2/3 pathway (Carnell et al., 2011; Lystad, 2025). Barrier assays also demonstrated that RNAseK RNAi flies exhibited strong Smurf positivity (**Fig. 4M-O**; **Fig. S3, A**).

### RNAseK is required for function of both major Malpighian tubule cell types

Malpighian tubules act as V□ATPase□powered analogues to the mammalian renal system, coordinating ion and water transport (Dow et al., 1994; Dow and Davies, 2001). Dual RNAseK depletion in principal and stellate cells caused severe abdominal enlargement (**Fig. 5A-B**), significantly increased abdominal width (**Fig. 5C**), and elevated water retention (**Fig. 5D**), representing systemic osmotic failure comparable to chloride□channel silencing (Cabrero et al., 2020). Knockdown tubules showed distorted epithelial architecture under polarised light (**Fig. 5E-F**) and intraluminal crystalline inclusions confirmed by Alizarin Red staining (**Fig. 5G-H**; **Fig. S5, D**), consistent with calcium precipitation documented in V□ATPase knockdown models (Allan et al., 2005; Dow et al., 2021).

Stellate cell specific RNAseK knockdown resulted in pronounced abdominal swelling (**Fig. 6A-C**) and elevated water content (**Fig. 6D**), demonstrating systemic fluid dysregulation. Microscopy showed severe smoothening of the tubule surface and depletion of ridged topology (**Fig. 6E-F)**, while Alizarin Red staining identified calcium deposits (**Fig. 6G-H**), signifying ionic imbalance (Dickson, 1965). These results reinforce the dependence of stellate□cell chloride and water channel function on V□ATPase□linked acidification (Cabrero et al., 2014; Dow and Davies, 2001).

Survival assays revealed reduced starvation tolerance upon stellate RNAseK knockdown (**Fig. 6I**) but elevated desiccation resistance (**Fig. 6J**), phenocopying fluid□retentive adaptations of V□ATPase or ClC□mutant flies (Cabrero et al., 2020). Mechanistically, this suggests that RNAseK may regulate coupling between apical acidification and vesicular trafficking, maintaining ClC□a and aquaporin localization required for diuresis (Perreira et al., 2015; Cabrero et al., 2014, 2020). To further explore how RNAseK disruption affects tubule epithelial tissue architecture, extensive imaging of control and RNAseK-knockdown flies, which revealed regularly spaced, actin-rich stellate cells in control tubules (**Fig. 6K**). In contrast, RNAseK RNAi markedly reduced stellate cell number and distorted their morphology (**Fig. 6L; Fig. S5, G)**. Overexpression of RNAseK specifically in stellate cells resulted in these cells appearing more prominent and distinct than in controls (**Fig. S5, G**), suggesting a role for RNAseK in promoting the proper shaping, spacing, and possibly the coordinated integration of stellate cells among principal cells during their mesenchymal-to-epithelial transition in tubulogenesis (Denholm et al., 2013). Quantitative analysis further demonstrated a significant decrease in stellate cell number in both anterior and posterior tubules upon RNAseK knockdown (**Fig. 6M**), reinforcing its importance in stellate cell invagination, distribution, and potentially long-term survival. Fluid-secretion measurements showed strong reduction under both unstimulated and CAPA-stimulated conditions (**Fig. 6N**), indicating compromised neuropeptide-driven diuresis (Davies et al., 2013).

Principal□cell RNAseK RNAi flies displayed abdominal swelling (**Fig. 7A-C**) and fluid accumulation (**Fig. 7D**; **Fig S6, A-D**), accompanied by cystic tubule morphology and white, non□birefringent crystalline deposits (**Fig. 7E-F**) indicative of calcium phosphate/xanthine crystals (Ghimire, 2019) and further supported by Alizarin Red staining (**Fig. 7G-H**). Transcriptional profiling (**Fig. 7I**) revealed upregulation of SPoCk and Crystallin, known calcium□responsive genes (Southall et al., 2006; Chung and Turney, 2017), but no change in Vha68□2, supporting a post□translational regulatory function of RNAseK.

Functionally, RNAseK–deficient flies exhibited reduced starvation (**Fig. 7J**) and desiccation tolerance (**Fig. 7K**), consistent with metabolic imbalance and impaired water excretion similar to LKR mutants with defective diuresis (Cannell et al., 2016). These results establish RNAseK as a potentially critical regulator of principal□cell calcium handling and fluid homeostasis.

### RNAseK colocalises with V-ATPase to maintain epithelial membrane structure

To assess RNAseK localisation, we generated an anti-RNAseK antibody. Efficacy was tested by comparing across RNAseK expression levels; the fluorescence intensity of the antibody staining increased upon genetic upregulation (**Fig. S7, C**) and decreased when expression was intentionally genetically reduced (**Fig. S7, D**), confirming that the detected signals were directly correlated with RNAseK abundance and not due to non-specific binding,

The V-ATPase B-subunit, Vha55, is localised apically in tubule cells and restores fluid transport upon transgenic rescue (Du et al., 2006). Within the tubule, RNAseK exhibited complete apical colocalization with Vha55 fluorescence (**Fig. 8A**; **Fig. S7, A&E**), supporting direct cooperative interplay (Perreira et al., 2015). Upon RNAseK depletion, the protein redistributed diffusely from apical to basal membranes (**Fig. 8B; Fig. S7, F**), mirroring polarity-collapse phenotypes reported for V-ATPase loss in mammalian epithelia (Bidaud-Meynard et al., 2022). This indicates that RNAseK and V-ATPase function as an integrated apical complex essential for epithelial polarity.

Further, to assess the impact of RNAseK downregulation on cellular arrangement and boundaries, Dlg — a known surface marker within the tubule (Dornan et al., 2023) — was immunostained. Control tubules marked well□defined cell boundaries (**Fig. 8C**), while RNAseK RNAi tubules displayed irregular cell distribution and arrangement (**Fig. 8D**). Since Dlg interacts with adducin and actin to maintain epithelial architecture and polarization (Woods et al., 1996; Bilder et al., 2000; Wang et al., 2011), defects upon RNAseK downregulation in the form of disrupted apical F-actin organization correlated with disrupted Dlg organisation (**Fig. 8G**; **Fig. S6, H**). RNAseK knockdown affects epithelial integrity and leads to fused apical□basal layers observed under magnified views (**Fig. 8G**, blue arrow), also affects brush□border integrity; disintegration (**Fig. 8H**; **Fig. S6, H**) confirming global epithelial disarray upon RNAseK depletion, potentially linking vesicular acidification failure to polarity complex instability.

## Discussion

RNAseK has recently emerged as a critical regulator of proton pump function in mammalian systems (Perreira et al., 2015; Makar et al., 2024). Here, we have provided computational and biophysical evidence that this role for RNAseK is broadly conserved across evolutionary history, and RNAseK remains a crucial epithelial determinant in the fruit fly *Drosophila melanogaster* (Allan et al., 2005; Mo et al., 2020).

At the sequence level, the functional elements of RNAseK are strongly conserved even in species as distant from the fly as humans and zebrafish, suggesting that these regions effect vital biological functions. One may hypothesize that this function is linked to the recent reports of RNAseK as a regulatory component of the V-ATPase complex (Perreira et al., 2015; Makar et al., 2024), especially given that we have located multiple dCLEAR cis-regulatory elements in close proximity with the RNAseK gene, suggesting a shared transcriptional program with the epithelial vha genes (Bouché et al., 2016).

Our molecular experimentation supports the hypothesis that RNAseK is crucial in the major epithelial tissues within the fruit fly – namely the midgut, hindgut, Malpighian tubules and salivary glands – in a fashion consistent with proton pump disruption. Indeed, the major biological disruptions caused by RNAseK manipulation can broadly be linked with those described in V-ATPase disruption, including loss of polarisation during embryogenesis (Davies et al., 1996; Wieczorek et al., 2009), disruption of gut polarity and acidification (Overend et al., 2016), reduced starvation resistance (Cannell et al., 2016), disruption of tubule transport systems (Cabrero et al., 2020), impaired tubular fluid secretion and hormone response (Davies et al., 2013), defective secretory granule maturation in salivary glands (Farkaš et al., 2015), and defective V-ATPase membrane localisation (Bidaud-Meynard et al., 2022).

The exquisite dosage sensitivity of RNAseK is illustrates that both reduction and elevation of its levels in principal cells abolish normal xanthine pigmentation of luminal concretions, yet only depletion elicits crystal precipitation and induction of calcium-responsive genes such as SPoCk and Crystallin (Southall et al., 2006; Chung and Turney, 2017). This bell-shaped response argues that RNAseK acts as a rheostat that calibrates vesicular or luminal pH range for orderly urate deposition; any deviation disrupts pigment incorporation while excess RNAseK apparently maintains sufficient acidification to prevent uncontrolled nucleation. The Smurf phenotype and dye leakage observed upon RNAseK knockdown provide functional evidence of a profound breach in paracellular barrier integrity that is similar to the chronic epithelial leakage characteristic of aged flies and of mutants in septate-junction components (Rera et al., 2012; Salazar et al., 2023; Dornan et al., 2023), suggesting that V-ATPase activity, maintained by RNAseK, is continuously required throughout adult life.

Likewise, the cortical F-actin collapse seen in the midgut upon RNAseK depletion mirrors phenotypes produced by disruption of the WASH–Arp2/3 actin-nucleation pathway (Carnell et al., 2011), raising the possibility that RNAseK supports V-ATPase-dependent retrieval or recycling of actin-regulatory complexes at the apical membrane, thereby linking proton-pump fidelity directly to cytoskeletal homeostasis in epithelia. Hindgut-specific RNAseK downregulation causes localized swelling immediately anterior to the tubule-ureter junction, accompanied by fluid backflow, retention, and white/black deposits just posterior to the ureter junction that occlude the lumen. This is explained by the potential loss of the strong electrogenic H□gradient normally generated by hindgut V-ATPase, which abolishes the driving force for K□/H□exchange and Cl□-driven water reabsorption, resulting in osmotic water retention and precipitation of undissolved waste (Shanbhag and Tripathi, 2005; Overend et al., 2016).

Finally, the ability of RNAseK depletion to trigger rapid basolateral mislocalisation of Vha55 and brush-border dissolution (Du et al., 2006; Dornan et al., 2023) identifies RNAseK as the possible apical tether for the holoenzyme in post-mitotic epithelia. As such, while we have not directly proven molecular interaction, we have gathered a bevy of evidence supporting the notion that RNAseK is absolutely required in epithelial tissues to support the normal functioning of the V-ATPase holoenzyme. Collectively, our findings propose RNAseK as a regulator of V-ATPase performance across epithelia. These findings are highly relevant not only in Drosophila but, given the high degree of functional RNAseK homology across evolutionary history, also have implications for mammalian physiology and medical science.

## Acknowledgements

We thank Dr Robin Beaven (University of Edinburgh), Dr Adam Dobson (University of Glasgow), and the Ilan Davies lab (University of Glasgow, Scotland, UK) for generously providing the Hindgut-GAL4/CyO; UAS-GFP/TM6c, Myo31df-GAL4; tub Gal80^TS^, and Da GAL4; tub Gal80^TS^ fly lines, respectively. Additionally, we thank the Dr. Alberto Sanz group (University of Glasgow) for their ongoing support and provision of the DAMs system utilised here. We also thank the University of Glasgow MVLS SRF Cellular Analysis facility for their expert advice on confocal microscopy.

## Data Availability

All code used in this analysis is available at the Figshare repository 10.6084/m9.figshare.31037749

---

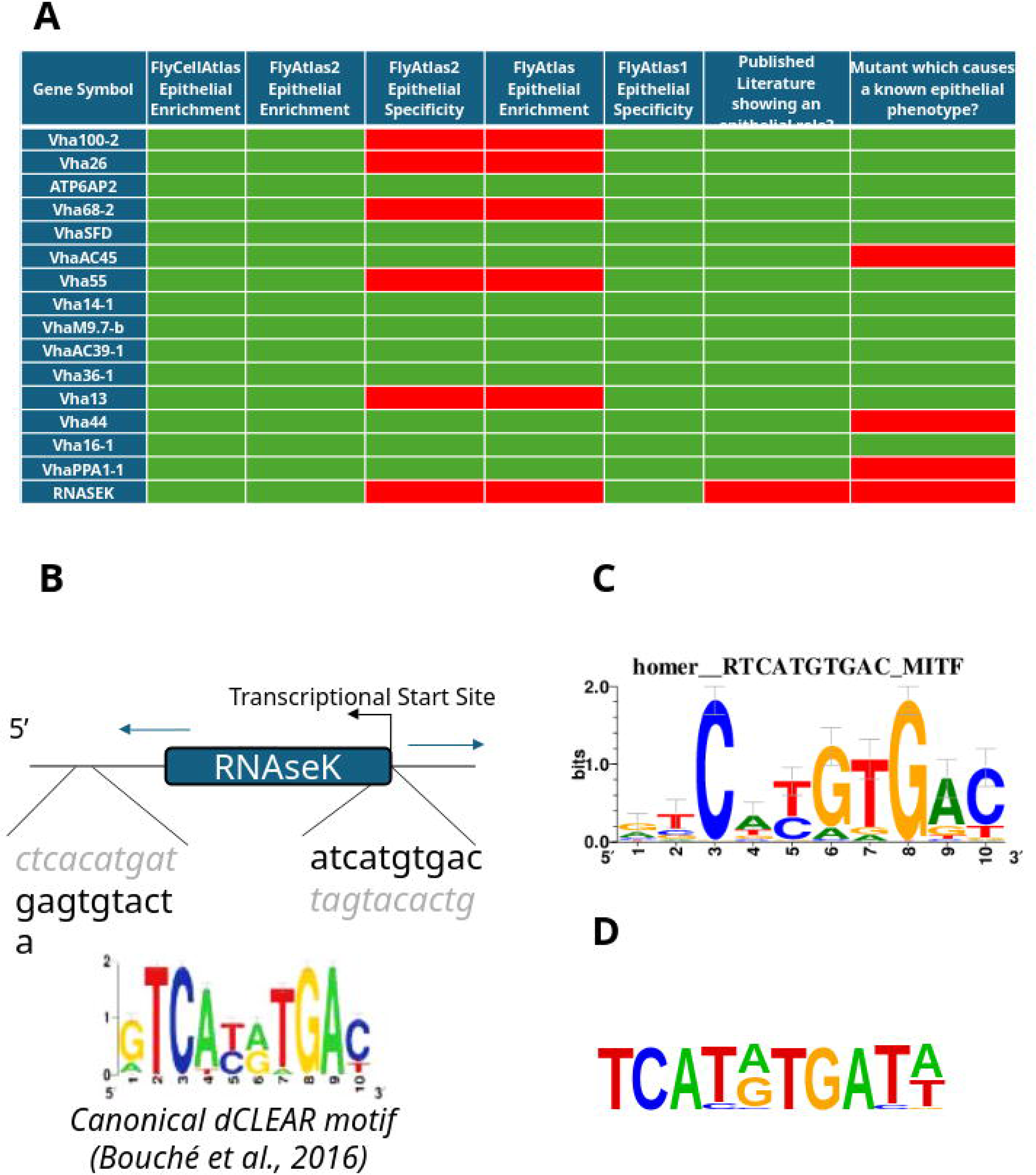

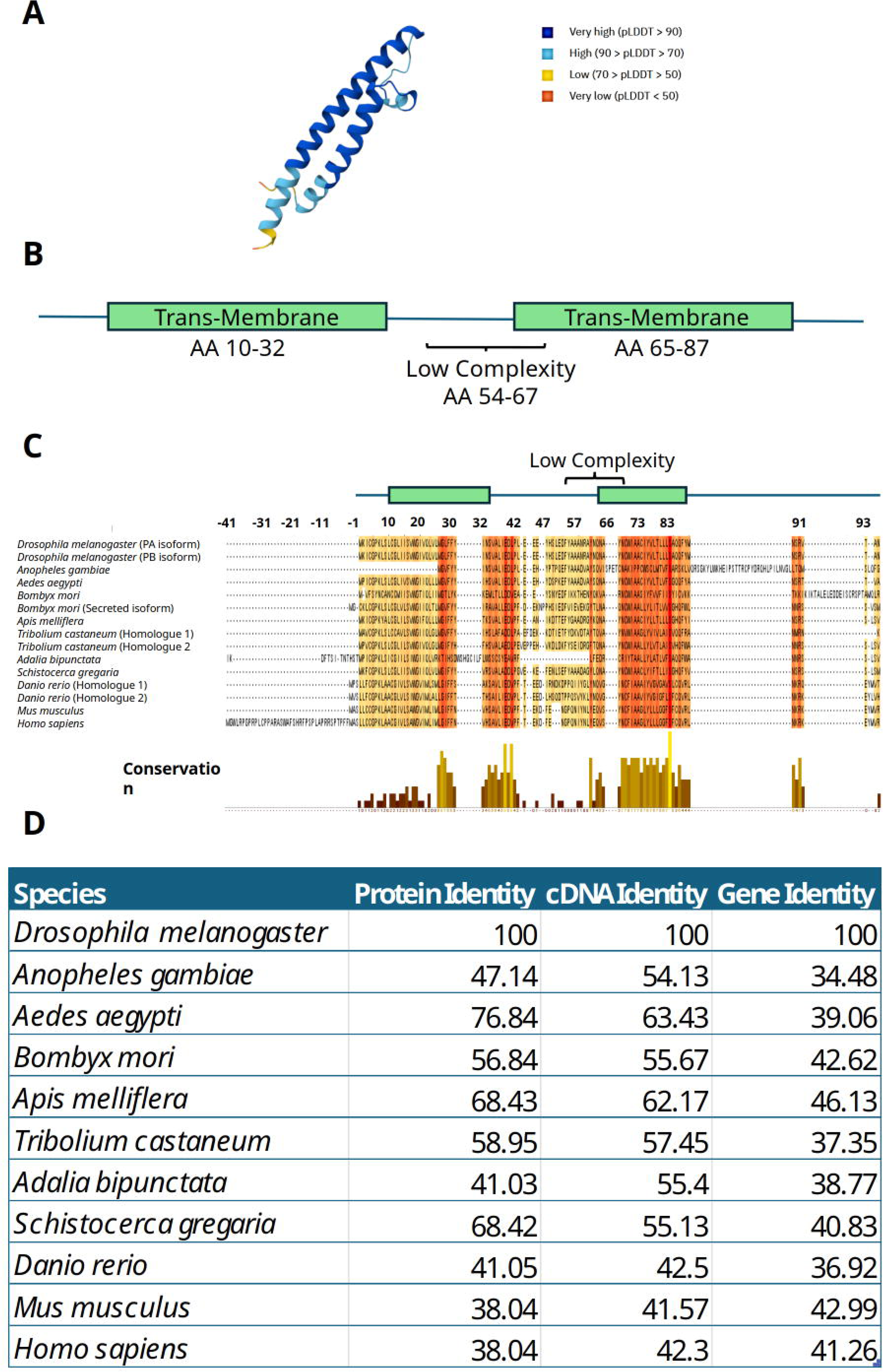

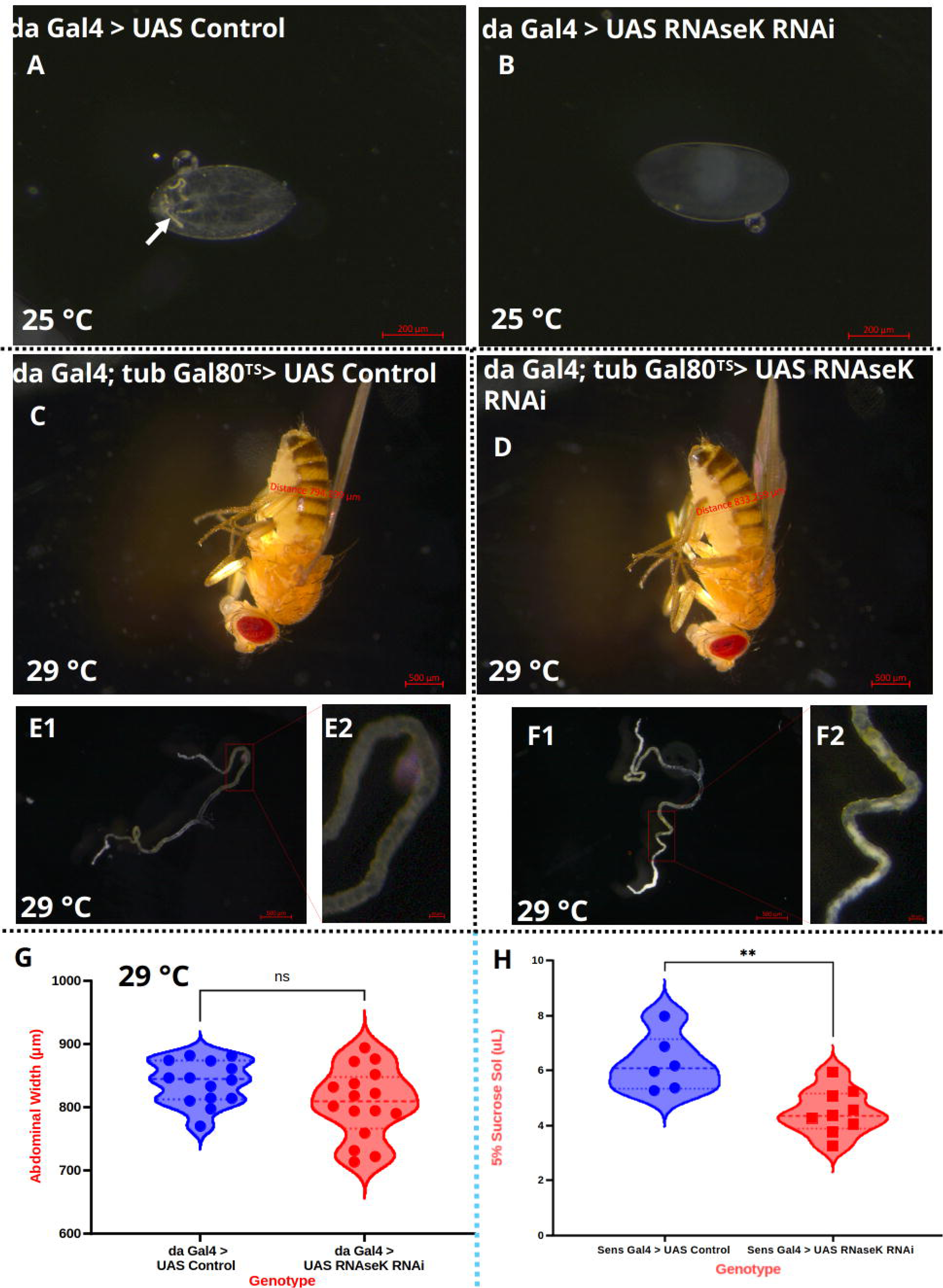

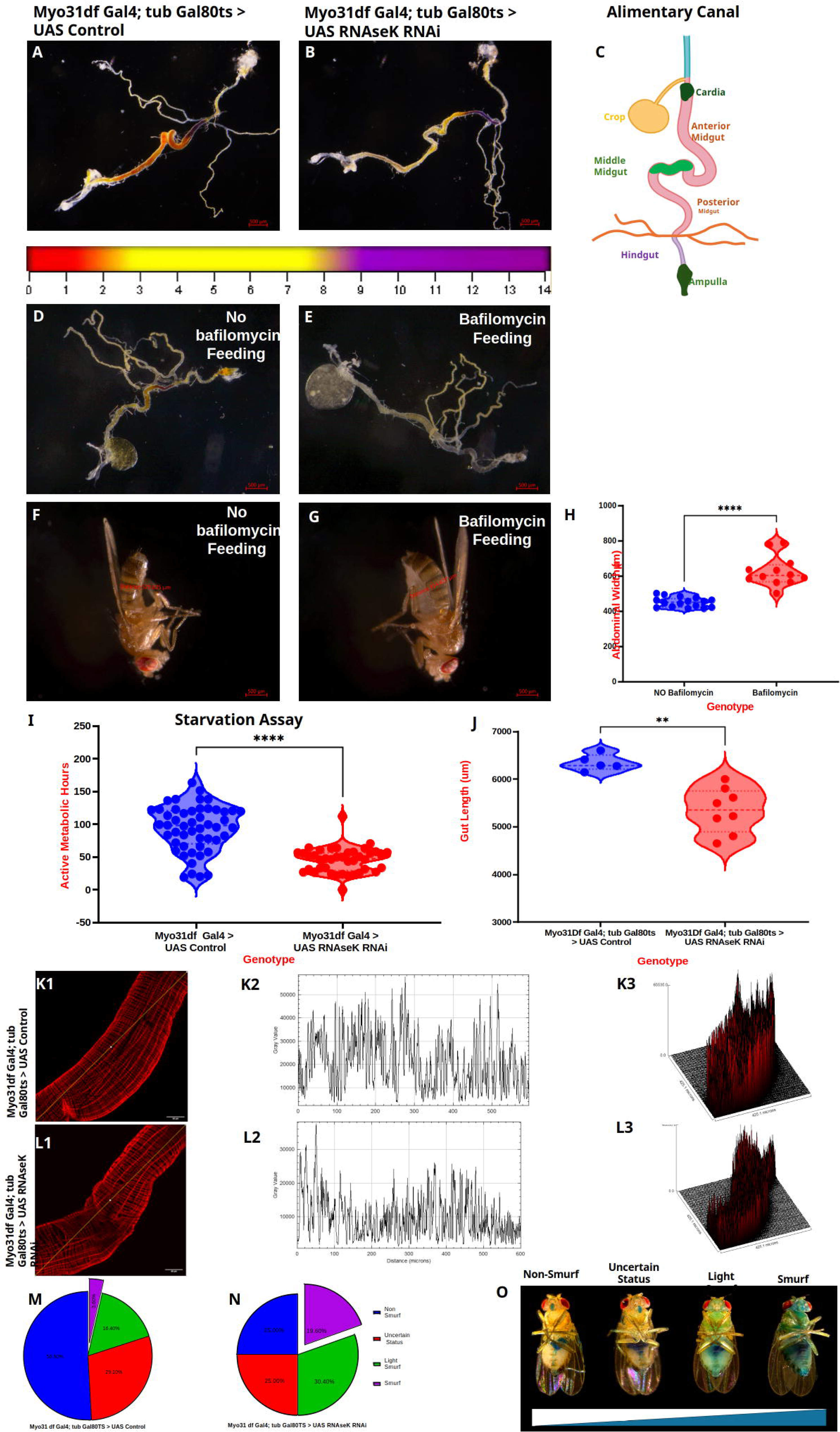

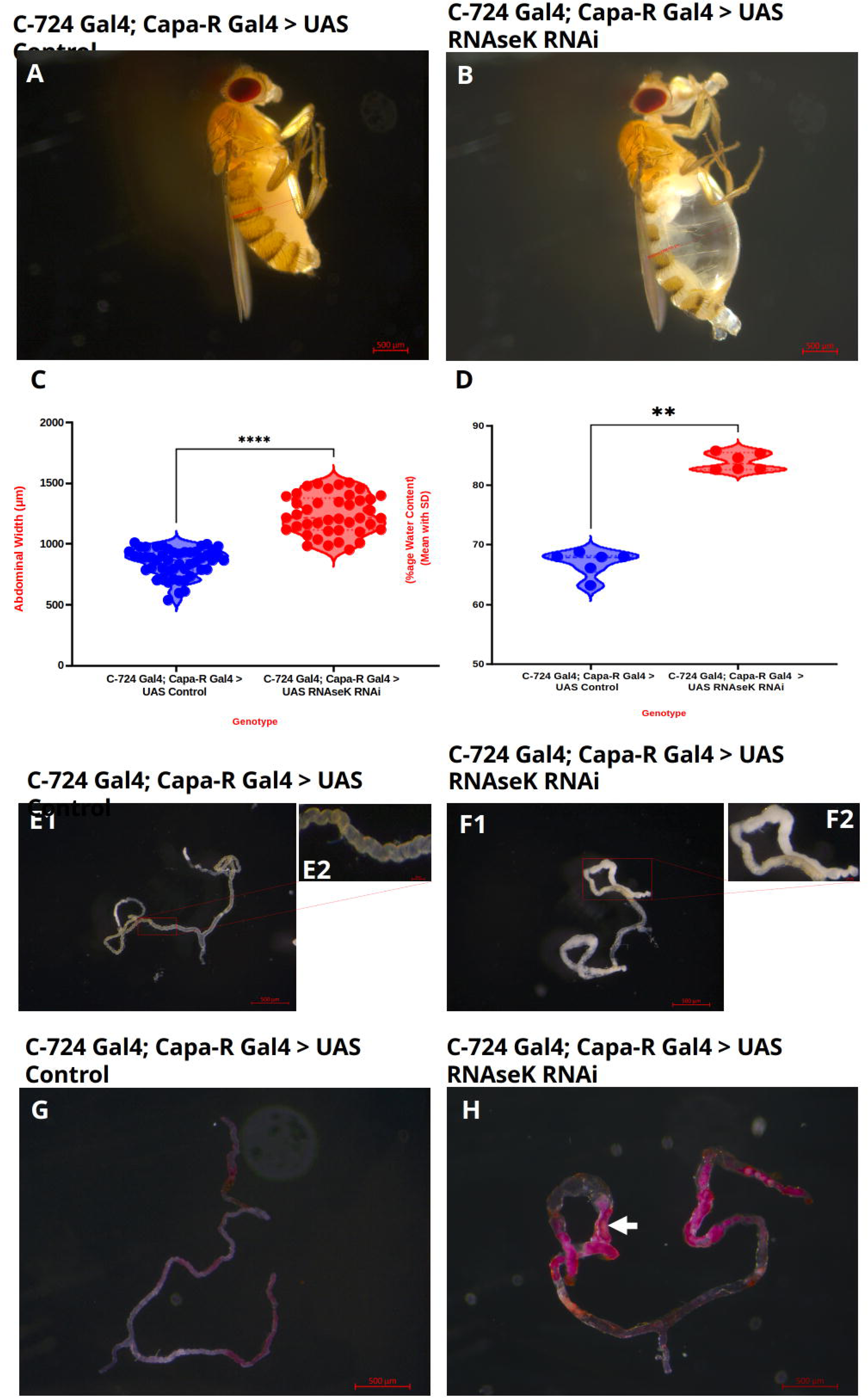

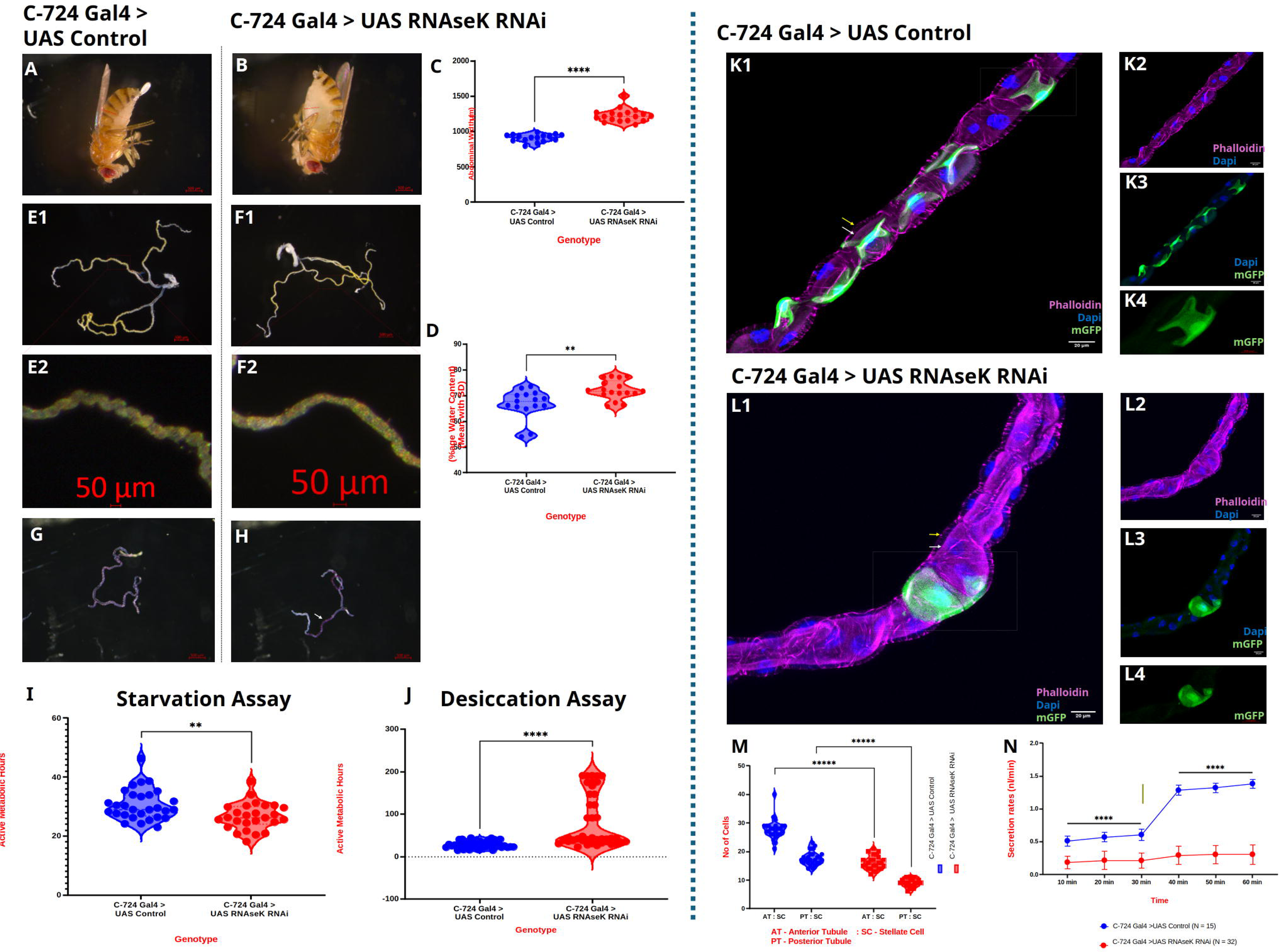

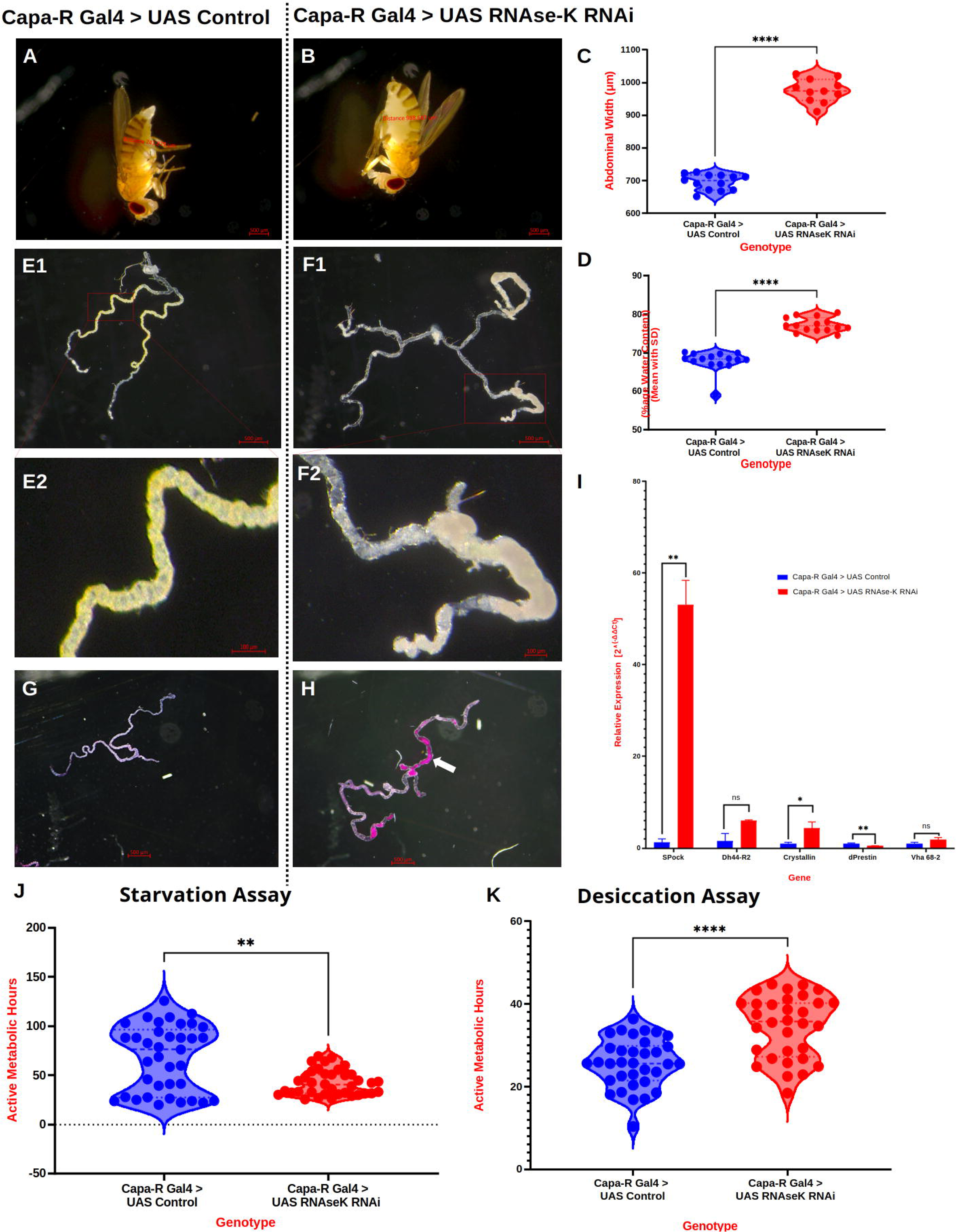

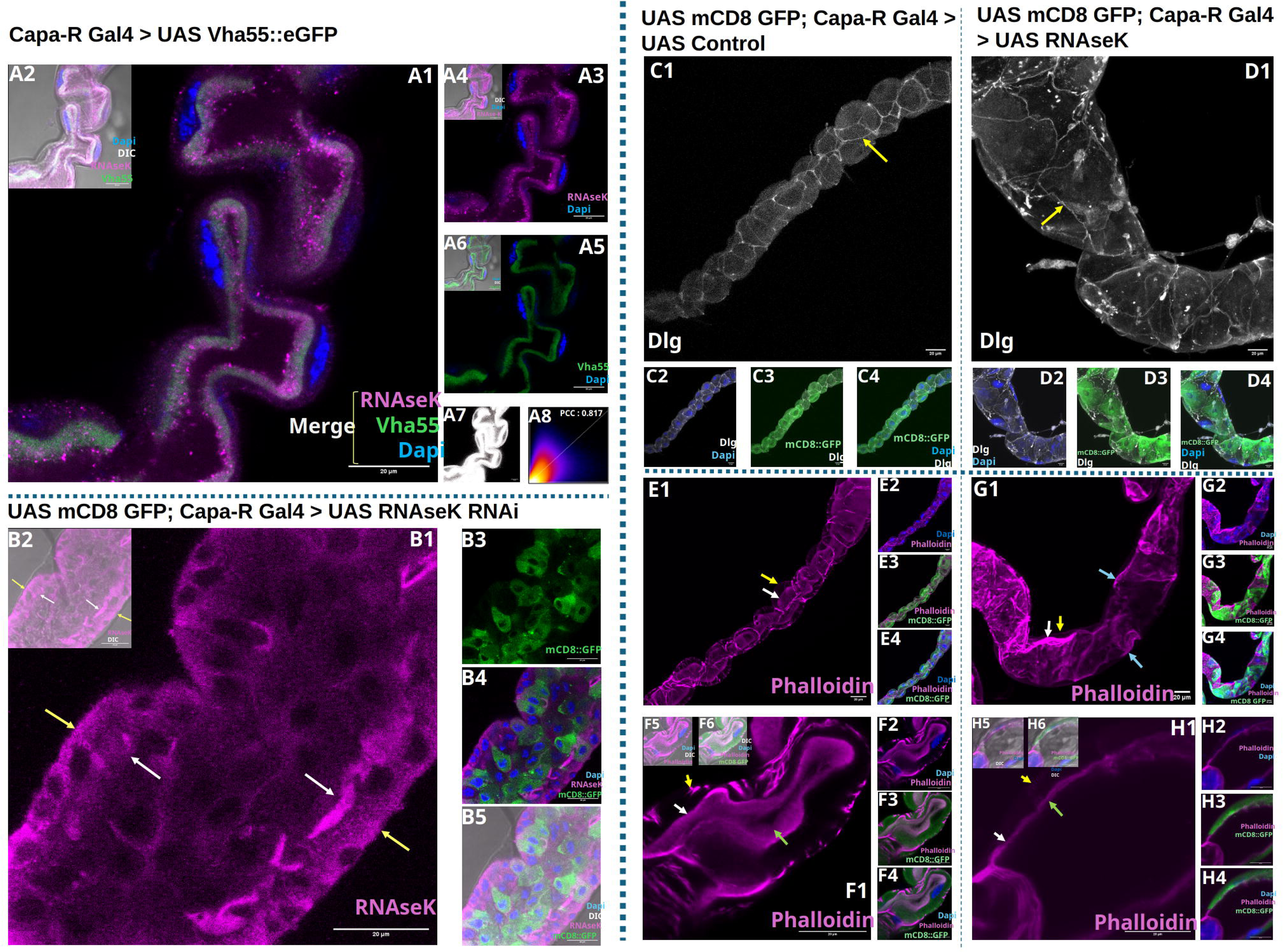

